# Low dose resveratrol promotes hypertrophy in wildtype skeletal muscle and reduces damage in skeletal muscle of exercised *mdx* mice

**DOI:** 10.1101/289587

**Authors:** KG Woodman, CA Coles, SL Toulson, M Knight, M McDonagh, SR Lamandé, JD White

## Abstract

Duchenne muscular dystrophy (DMD) is a progressive and fatal neuromuscular disorder for which there is no treatment. Therapies to restore dystrophin deficiency are not ready for clinical use and long-term efficiency is yet to be established. Therefore, there is a need to develop alternative strategies to treat DMD. Resveratrol is a nutraceutical with anti-inflammatory properties and previous studies have shown that high doses can benefit *mdx* mice. We treated 4-week-old *mdx* and wildtype mice with low-dose resveratrol (5mg/kg bodyweight/day) for 15 weeks. A voluntary exercise protocol was added to test if low dose resveratrol could reduce exercise-induced damage. We showed that resveratrol promoted skeletal muscle hypertrophy in the wildtype mice. There was no change in markers of pathology in the *mdx* mice; however, the low-dose resveratrol reduced exercised induced damage. Gene expression of immune cell markers such as CD86, CD163 and PCNA was reduced; however signalling targets associated with resveratrol’s mechanism of action of action including SIRT1 and NF-κB were unchanged. In conclusion, low-dose resveratrol was not effective in reducing disease pathology; however, its ability to promote hypertrophy in wildtype skeletal muscle could have direct applications to the livestock industry or in sports medicine.

## Introduction

Duchenne muscular dystrophy (DMD) is a progressive neuromuscular disorder, that arises from mutations in the dystrophin gene (Koenig et al., 1988) and leads to the absence or severe deficiency of the dystrophin protein (Hoffman et al., 1987). In DMD corticosteroids are used to prolong ambulation and maintain respiratory health; however, they are associated with a number of adverse side effects (Biggar et al., 2004; Bushby et al., 2004; Hoffman et al., 1987). Many types of therapeutics are in development for DMD and include gene correction strategies, exon skipping or utrophin upregulation, however, these therapies are some time away from being implemented into a clinical setting and their long-term efficiency is yet to be evaluated. Therefore, there is a need to develop alternative strategies to treat DMD which could be implemented in the short term or in the longer term as an adjunct to gene correction strategies.

Nutraceutical usage has been increasingly popular in recent years and this trend has not escaped the DMD community (reviewed in Woodman et al., 2016). While nutraceuticals won’t correct the genetic defect in DMD, many have either anti-inflammatory or anti-oxidant properties which could potentially treat the secondary processes such as chronic inflammation which occur in DMD patients. Given that many nutraceuticals are already in use or available over the counter they can be readily implemented into a clinical setting.

Resveratrol (3,5,4-trihydroxy-trans-stilbene), a polyphenol found in grapes, is a nutraceutical that affects pathways such as carcinogen metabolism, cell proliferation, inflammation, cell cycle regulation and apoptosis (reviewed in Woodman et al 2016). Many of resveratrol’s actions occur due to activation of sirtuin 1 (Sirt 1). Resveratrol allosterically binds to the N terminus of Sirt1 thus activating it (Hubbard et al., 2013). Sirt1 subsequent deacetylates a range of downstream signaling targets in a variety of tissue types.

A series of studies have considered the effect of resveratrol on the viability and differentiations of the C2C12 cell line (Kaminski et al., 2012; Montesano et al., 2013; Wang et al., 2016). While these studies used varying concentrations of resveratrol the key findings included decreased cell viability (Kaminski et al., 2012) and increased myoblast elongation and differentiation; this was associated with increased expression of the myogenic regulatory factors (MRFs) *MyoD and myogenin* (Kaminski et al., 2012). The effect of resveratrol on the expression of MRFs differs depending on whether cells are proliferating or differentiating; resveratrol increases expression of *Myf5* and *MyoD* during the proliferation phase and *myogenin* and MyHC during the differentiation phase (Montesano et al., 2013). Studies using the Sirt1 inhibitor, nicotinamide, directly link the effect of resveratrol on C2C12 cells with Sirt1 expression; proliferation was decreased with nicotinamide treatment (Wang et al., 2016).

The *in vitro* effects of resveratrol on myogenic behavior led to a series of studies in the *mdx* mouse model of DMD. The studies used different resveratrol doses and assessed different outcomes. Delivery of 100mg/kg/day for eight weeks resulted in a significant reduction in bodyweight and *EDL*, soleus and *TA* muscle weights (Selsby et al., 2012). In the same study fatigue resistance was increased approximately 20% in the soleus muscle but the *EDL* and *soleus* muscles were protected from contraction-induced injury. A high mortality rate has been observed when resveratrol has been used at higher doses [400 mg/kg/day]. When resveratrol is delivered to mdx mice via oral gavage for 10 days at 10, 100 or 500 mg/kg only the 100 mg/kg dose resulted in a significant increase in Sirt1 gene expression (Gordon et al., 2013). This was associated with decreased immune cell infiltration in the *gastrocnemius* muscle but no reduction in gene expression of the pro-inflammatory cytokine *TNF-a.* In a later study 100mg/kg resveratrol treatment did not improve grip strength or fatigue in the *mdx* mice but produced a significant improvement in rotarod performance (Gordon et al., 2014).

These studies have demonstrated resveratrol treatment has positive effects in dystrophic muscle. We used much lower doses of resveratrol (5mg/kg)in rations for growing lambs and showed significant changes in the growth of lean tissue with no increase in fat (manuscript in preparation). Here, we assesses a dose at the lower end of what has been previously tested, over a longer time period, employing a comprehensive analysis of muscle pathology and incorporating voluntary exercise into the experimental protocol to increase damage in dystrophic tissue and better reflect the human condition.

## Methods

All reagents were purchased from Sigma Aldrich (St Louis, Missouri, USA) unless otherwise specified.

### Animal Ethics

All animal experiments were approved by the University of Melbourne Animal Ethics Committee (AEC) and the Murdoch Children’s Research Institute AEC. Animals were cared for according to the ‘Australian Code of Practice for the Care of Animals for Scientific Purposes’ published by the National Health and Medical Research Council (NHMRC) Australia (National Health and Medical Research Council, 2013) and were housed under a 12 hour light/dark cycle with food and water provided *ad libitum*.

### Preparation of Nutraceutical Diets

Resveratrol was obtained from Sigma Aldrich and was sent to Specialty Feeds for preparation (Glen Forrest, Western Australia, Australia). The dose was determined assuming the standard mouse weighs 30g and consumes 5g feed daily. Resveratrol was added to standard mouse chow to produce a dosage of 5mg/kg bodyweight/day. The control diet consisted of the standard mouse chow, from the same manufacturer, without any additives.

### Trial Design

Male C57BL10/ScSn (wildtype) and C57Bl10/mdx (mdx) mice were obtained from the Animal Resources Centre (Perth, Western Australia, Australia). At 4 weeks of age, 12 mice received either a control diet or a diet containing resveratrol (5mg/kg bodyweight/day) for 12 weeks. After 12 weeks the groups were halved, with one half allowed voluntary exercise (exercise group) and the other half housed normally (sedentary group). Mice were exercised for three weeks, including a one week acclimatization period where multiple mice were housed in the cage with the wheel to become accustomed to it, followed by a two week exercise period where mice were housed in separate cages containing voluntary running wheels (Smythe and White, 2011). The mice continued to receive their specified diet over the exercise period. On completion of the feed trial blood was obtained via cardiac puncture and the quadriceps muscles were dissected out and snap frozen in liquid nitrogen. Once frozen the muscles were stored at -80°C for RNA and protein extractions. The contralateral muscles from the remaining hindlimb were mounted in 5% tragacanth (w/v) and frozen in liquid nitrogen cooled isopentane and stored at -80°C for histological analyses.

### Histological Sample Preparation

The quadriceps muscle is more susceptible to exercised-induced damage than other muscles (Archer et al., 2006; Grounds et al., 2008; Hodgetts et al., 2006). Therefore; the *quadriceps* muscle was used to analyse the effects of the diets on muscle morphology and histopathology. The frozen *quadriceps* muscle was transversely cryosectioned on a Leica CM3050S cryostat (Leica Biosystems, Vista, California, USA) at a thickness of 10μm. Sections were mounted onto Superfrost Plus microscope slides (Menzel-Glaser, Brunswick, Germany) and stored at -20°C until required.

### Immunohistochemistry

Frozen sections were thawed, and a Pap pen (ProSciTech, Kirwan, Queensland, Australia) was used to draw a boundary between each section prior to rehydration in 1xPBS for 30 minutes. Sections were blocked in 10% (v/v) donkey serum (Millipore, Billerica, Massachusetts, USA) in wash buffer (0.1% Tween, 0.5% BSA in 1xPBS) for one hour, and then incubated with the primary antibody (laminin α2) diluted in wash buffer (1:200) overnight at 4°C. The sections were washed 3 times in wash buffer and then incubated with fluorescent secondary antibodies (AlexaFluor594⍰donkey anti-rat IgG⍰1:250 and donkey anti-mouse IgG 1:250 in wash buffer) in the dark for 90 minutes. Sections were then washed 3 times with wash buffer and stained with 1μg/μl Hoechst (Life Technologies, Carlsbad, California, USA) diluted to 1:1000 in 1xPBS for one minute before mounting with polyvinyl alcohol with glass coverslips. Sections were imaged on a Zeiss Axio Imager M1 upright fluorescent microscope with an AxioCam MRm camera running AxioVision software V4.8.2.0 (Carl Zeiss, Oberkochen, Germany).

### Measures of Histopathology

In all histological and morphometric assessments the total cross-section of the *quadriceps* was analysed, therefore a minimum of 2500 myofibres per animal were assessed. The treatment groups for these experiments were blinded to prevent any experimenter bias.

### Minimum Feret’s Diameter

Sections stained with laminin α2 were used to determine myofibre diameter. Images were analysed in Image J version 1.48G (U. S. National Institutes of Health, Bethesda, Maryland, USA). The image threshold was set and minimum Feret’s diameter was calculated according to the Treat-NMD standard operating procedure “Quantitative determination of muscle fibre diameter” (http://www.treat-nmd.eu/downloads/file/sops/dmd/MDX/DMD_M.1.2.001.pdf).

### Central Nucleation

Images were opened in Image J and the number of myofibres per image was calculated. The number of fibres with peripheral nuclei were counted manually and subtracted from the total number of myofibres, thus resulting in the number of myofibres with centrally located nuclei. The number of myofibres with central nuclei per quadriceps cross section was expressed as a percentage and averaged over each treatment group.

### Measurement of leaky myofibres

To determine the percentage of leaky myofibres in the *quadriceps*, the muscle was transversely sectioned and stained with laminin α2 to stain the basement membrane of the myofibres and IgG, which enters damaged myofibres with weakened sarcolemma. IgG positive myofibres over the muscle cross-section were counted manually and expressed as a percentage of the total myofibre number.

### Measurement of skeletal muscle necrosis

To assess the amount of necrosis present in the *quadriceps* muscle the transverse sections were stained with haematoxylin and eosin (H&E). Using Image J the total area of the muscle cross-section was calculated. Areas of necrosis were defined based on the presence of infiltrating inflammatory cells and areas of degenerating myofibres with fragmented sarcoplasm according to Treat-NMD standard operating procedure (http://www.treat-nmd.eu/downloads/file/sops/dmd/MDX/DMD_M.1.2.007.pdf). The amount of necrosis was expressed as a percentage of the total quadriceps area.

### Creatine Kinase assay

Blood was centrifuged at 12,000g for 15 minutes to separate out the serum. Serum was aliquoted into sterile tubes and stored at -80°C until required.

To measure CK activity, the serum was thawed and 5μl was aliquoted into a 96 well plate in triplicate, followed by 100μl of CK-NAC reagent (Thermo Scientific, Waltham, Massachusetts, USA). The change in absorbance was recorded at 340nm over three minutes (measured in 20 second intervals) at 37°C using a Paradigm Detection Platform (Beckman Coulter, Brea, California, USA) (Hunt et al., 2011).

### RNA extraction, cDNA synthesis, qPCR and oligonucleotide primer design

RNA was extracted from the snap frozen *quadriceps* by adding 1mL of TriReagent and homogenised using a T10 Basic S5 hand held homogeniser (IKA, Rawang, Selangor, Malaysia). To this, 100μl of chloroform replacement; 1-bromo-3-chloropropane was added, tubes were thoroughly vortexed and incubated at room temperature for 10 minutes. The RNA-containing aqueous phase was carefully removed and added to an equal volume of SV binding buffer (containing one part 90% ethanol and one part SV lysis buffer from SV Total RNA Isolation System (Promega, Madison, Wisconsin, USA)). The remaining procedure was as per the manufacter’s instructions. Concentration and purity of the RNA was determined using a NanoDrop1000 (NanoDrop, Wilmington, Delaware, USA) spectrophotometer at both 260 and 280nm.

For each reverse transcription reaction 1μg of RNA was added to a sterile tube. To each reaction 1.25μl of 10mM deoxyribonucleotides (dNTPs) and 1μL of 0.5mg/mL random oligo-hexamers (Promega, Madison, Wisconsin, USA) was added the reaction was performed as per the manufacturer’s instructions.

Each qPCR reaction contained 0.5μl of cDNA, 4μl of nuclease free water, 0.25μl of both forward and reverse primers (at a stock concentration of 20μM) and 5μl of ‘Go Taq Sybr Green’ qPCR master mix (Promega, Madison, Wisconsin, USA). All qPCR reactions were performed in white, 384 well qPCR plates (Roche Applied Science, Basel, Switzerland) using a LightCycler480 (Roche Applied Bioscience, Basel, Switzerland) calibrated to detect the Sybr Green chemistry (Hunt et al., 2011).

*Hprt* was chosen as the housekeeping gene as we have previously demonstrated it was the least variable across all mdx and wildtype muscle samples compared to other housekeepers using the ‘‘Bestkeeper’’ excel-based tool (Pfaffl, 2004). Primers used for quantitative PCR (qPCR) were designed as previously described (Hunt et al., 2011)and are presented in Table 1.

**Table 1.**
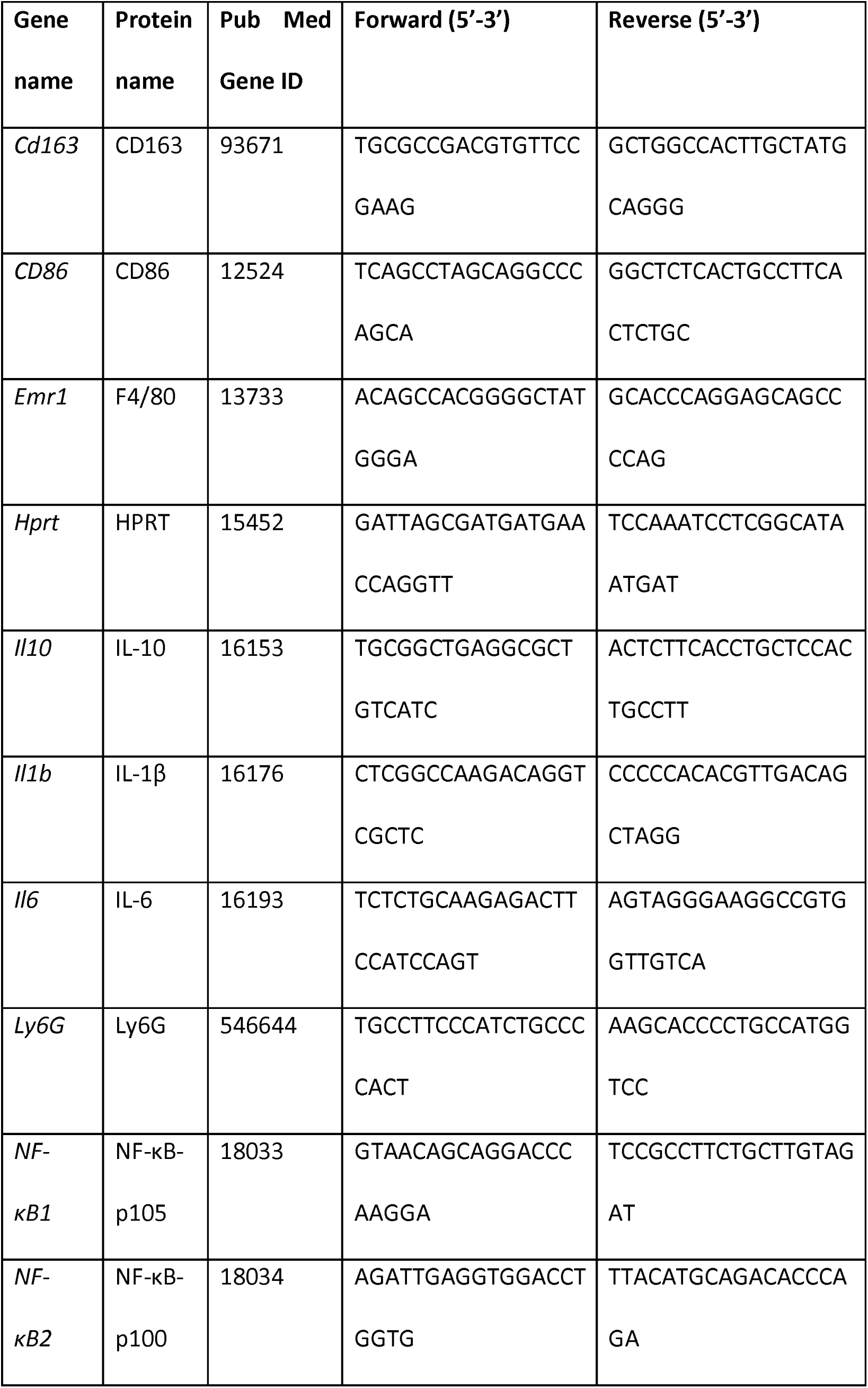
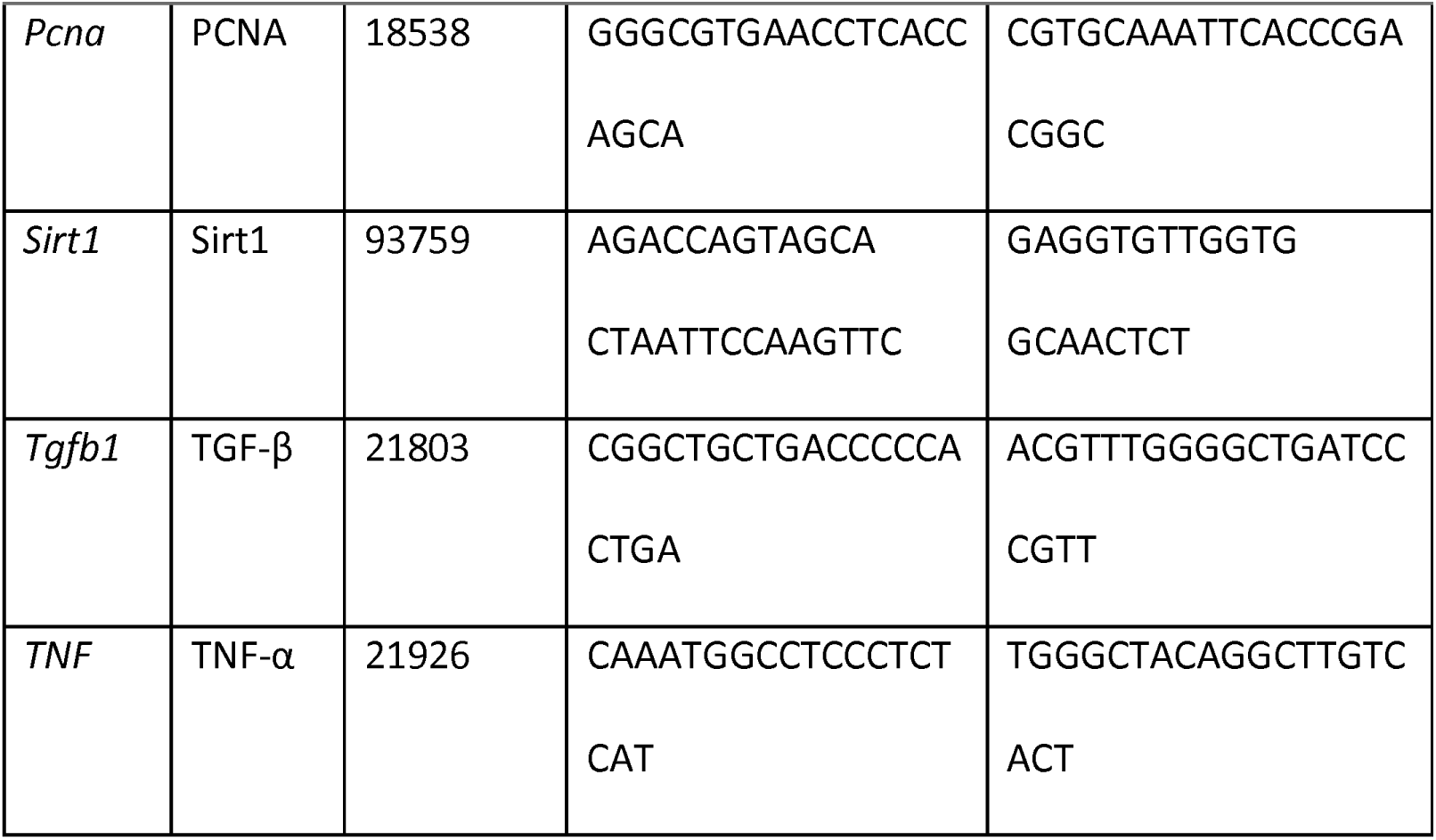
Primer sequences used for qPCR.

### Statistical Analyses

Where data was normally distributed GraphPad Prism 5 (GraphPad Software, La Jolla, California, USA) was used to assess statistical significance. Where there was a direct comparison between two groups or two data points, a Student’s t-test was used. For analysing multiple groups the statistical package Genstat (14^th^ Edition) (VSN International, Hemel Hempstead, Hertfordshire, United Kingdom) was used. A one-way analysis of variance (ANOVA), with Dunnet’s or Bonferroni post-hoc test was used when all treatments were compared to a single control or all treatments compared to each other respectively. qPCR data was assessed using a non-parametric Mann-Whitney U test in GraphPad. (Pfaffl, 2004).

## Results

### Effect of resveratrol administration on myofibre diameter in wildtype and mdx quadriceps muscle

The average myofibre diameter was larger in the resveratrol treated wildtype mice compared to the mice on the control diet (p<0.001) (Figure 1). In resveratrol treated wildtype mice there was a shift to larger myofibre diameters with increase in larger myofibres from 50-70μm in diameter when compared to the wildtype mice on the control diet (p<0.05) (Figure 1d). There was no difference in the average myofibre diameter in mdx mice between the resveratrol treatment and the control group (Figure 2), and no change in the myofibre size distribution in *mdx* mice treated with resveratrol (Figure 2d).

**Figure 1.**
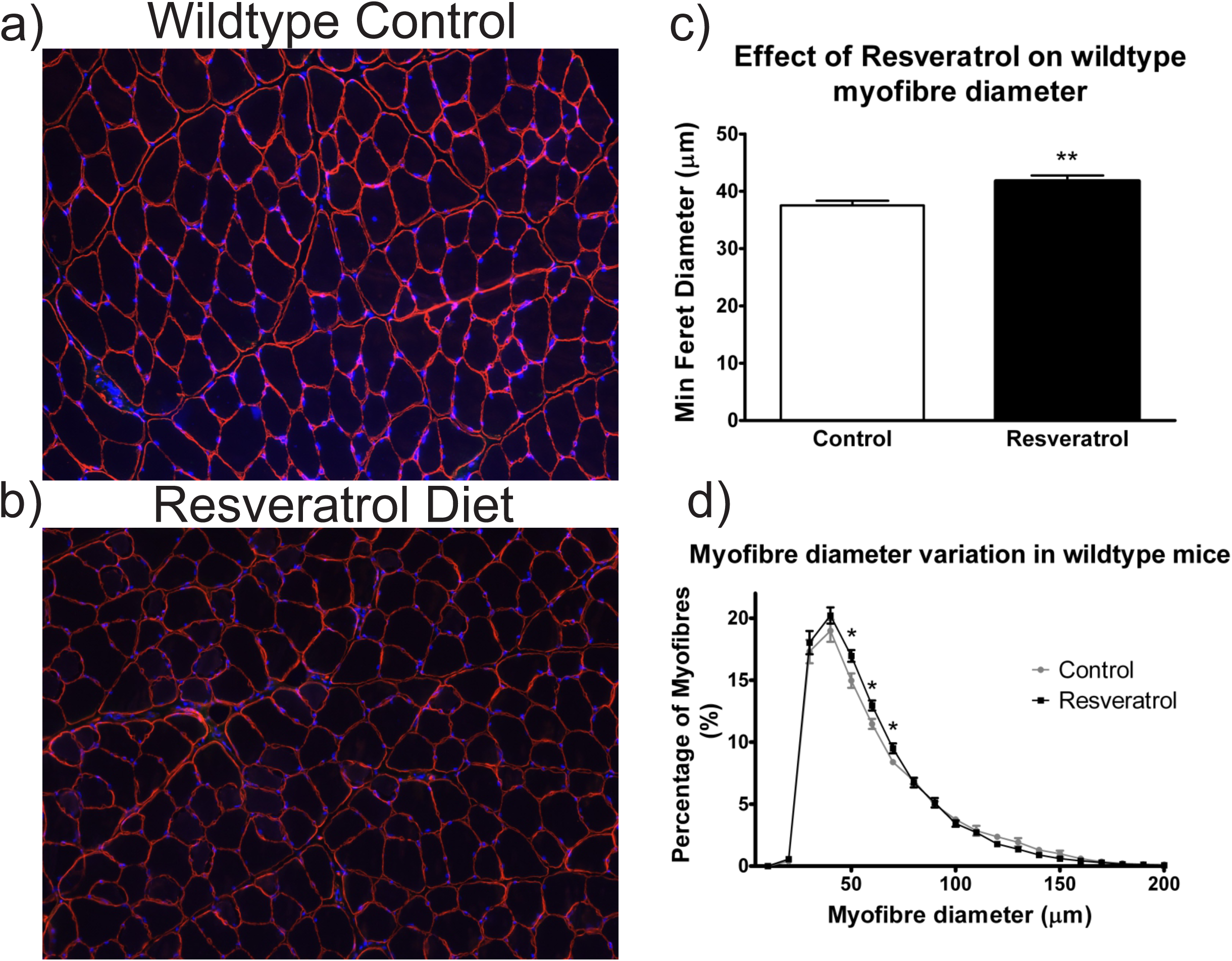
Resveratrol treatment increases myofibre diameter in wildtype *quadriceps*. Frozen *quadriceps* sections were stained with laminin α2 (red) which outlines the individual myofibres and Hoechst (blue) which stains the nuclei a) Representative image of a stained quadriceps from a mouse on the control diet. b) Representative image of a resveratrol treated mouse *quadricep*s. c) The average myofibre diameter within the *quadriceps* muscle is significantly larger in the resveratrol treated cohort in comparison to the control cohort. d) The myofibres were grouped into 10μm intervals and the proportion of myofibres in each interval was plotted in a histogram. There were more myofibres measuring between 50-70μm in the resveratrol treated group in comparison to the control group.. White scale bar indicates 200μm. Graphs show mean ± SEM. * indicates significance of p<0.05, ** indicates significance of p<0.001 (n=6).

**Figure 2.**
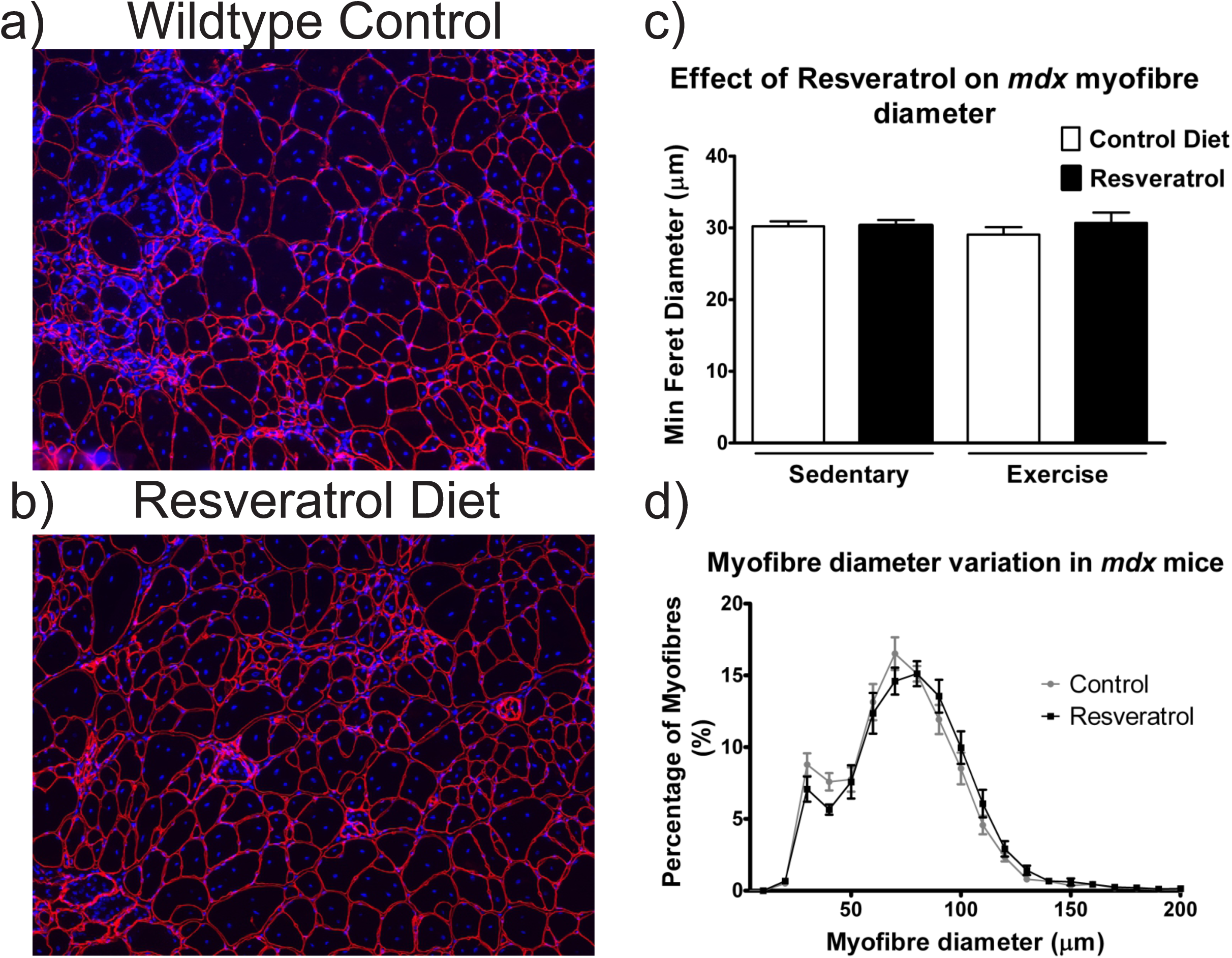
Resveratrol treatment does not affect myofibre diameter in *mdx quadriceps*. Frozen quadriceps sections were stained with laminin α2 (red) which outlines the individual myofibres and Hoechst (blue) which stains the nuclei a) Representative image of a stained *quadriceps* from a mouse on the control diet. b) Representative image of a resveratrol treated mouse *quadriceps*. c) The average myofibre diameter within the *quadriceps* muscle is unchanged with resveratrol treatment when compared to the controls. c) The myofibres were grouped into 10μm intervals and the proportion of myofibres in each interval was graphed. White scale bars represent 200μm. Graphs show mean ± SEM (n=6).

### Resveratrol treatment does not affect markers of muscle damage in non-exercised mdx mice

To determine if resveratrol treatment reduces dystrophic pathology in the *quadriceps* of the *mdx* mice, transverse frozen sections were stained with laminin α2, Hoechst and IgG (Figure 3 a-d). In mdx mice on the control diet voluntary exercise did not result in a significant increase in the percentage of fibers positive for IgG (Fig 3f). There was no significant difference in the percentage of myofibres staining positive for IgG with the resveratrol treatment in either the sedentary or exercised groups compared with mdx on the control diet (Figure 3e). Serum CK activity was not reduced with resveratrol treatment in either the sedentary or exercise groups when compared to the respective control diet cohorts (Figure 3f).

**Figure 3.**
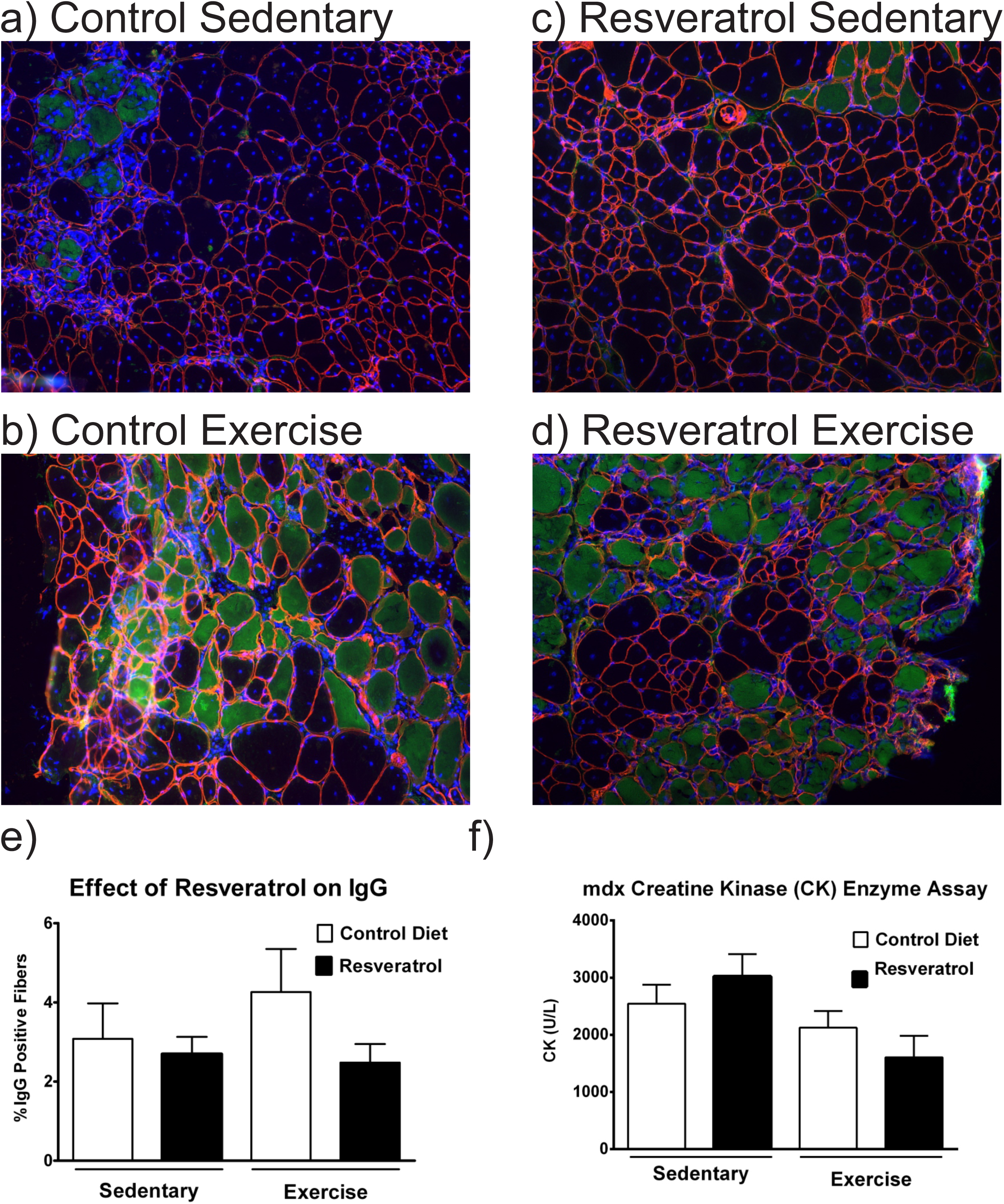
Resveratrol administration does not reduce the percentage of IgG fibres permeable in *mdx* mice. *Quadriceps* muscles were stained with laminin α2 (red) which outlines the myofibres, nuclei are stained in blue (Hoechst) and IgG in green which enters fibres with impaired sarcolemma. Representative image of an *mdx* control diet sedentary *quadriceps* (a), a control diet exercise mdx quadriceps (b), a resveratrol treated sedentary *quadriceps* (c) and a resveratrol treated exercise *mdx quadriceps* (d). e) Quantitative assessment of the *quadriceps* muscle shows the percentage of damaged, IgG positive fibres is unchanged with resveratrol treatment in both exercise and sedentary groups. f) Serum CK activity was similar in all cohorts. White scale bars represent 200μm. Graphs show mean ± SEM (n=6).

### Exercise-induced necrosis is reduced in mdx mice with resveratrol treatment

To determine if resveratrol administration reduced the amount of skeletal muscle necrosis present in the *mdx quadriceps* muscle the frozen transverse sections were stained with haematoxylin and eosin (representative images in Figure 4 a-d) and areas of necrosis were quantitated. The resveratrol treatment did not significantly reduce necrosis in the *quadriceps* of the sedentary *mdx* mice when compared to the control diet cohort (Figure 4e). The voluntary exercise protocol resulted in a significant increase in necrosis compared to the control sedentary *mdx quadriceps* (Figure 4e). This increased necrosis associated with the exercise protocol did not occur in the resveratrol treated *mdx* mice, indicating that the resveratrol treatment protected the muscle from voluntary exercise induced damage (Figure 4e).

**Figure 4.**
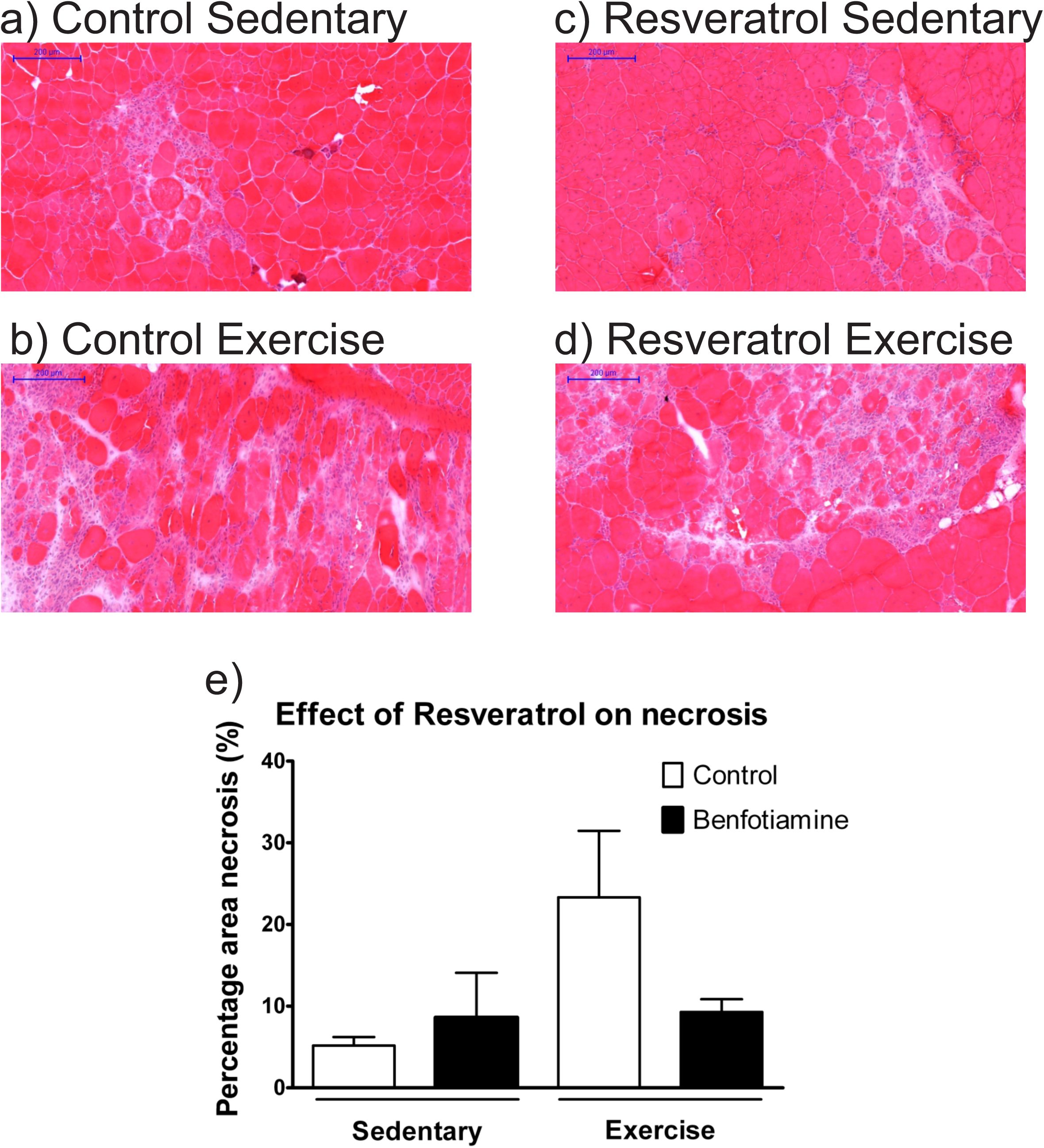
Resveratrol treatment prevents exercise induced muscle damage in *mdx quadriceps*. Transverse *quadriceps* sections were stained with haemotoxylin and eosin to illustrate muscle architecture, areas of inflammation and fibrosis. a) Representative image of an *mdx* control diet sedentary *quadriceps* (a), an mdx control diet exercise *quadriceps* (b), a resveratrol treated sedentary *mdx quadriceps* (c) and an mdx resveratrol treated exercised *quadriceps* (d). e) Necrosis was quantified and expressed as a percentage of the total *quadriceps* area. Necrosis was unchanged with resveratrol treatment in the sedentary mice compared to the control cohort. The reduced necrosis in the resveratrol treated exercise cohort was not significant. In the control diet mice there is a significant increase in necrosis with voluntary exercise. This effect is not observed in the resveratrol treated mice. White scale bars represent 200μm. Graphs show mean ± SEM. * indicates significance of p<0.05 (n=6).

### Resveratrol administration decreases expression of immune cell markers

The anti-inflammatory potential of resveratrol was investigated in *mdx* mice. Gene expression of immune cell markers in the *mdx quadriceps* muscle was assessed via qPCR. Expression of the pan-macrophage marker F4/80 (*Emr1)* was not changed by resveratrol treatment in either the sedentary or the exercised cohort when compared to the respective controls (Figure 5a). There was also no difference in expression of the neutrophil marker lymphocyte antigen 6 complex, locus G (Ly6G) gene with resveratrol treatment in either sedentary or exercise groups (Figure 5b). Expression of *Cd86* (Figure 5c), a marker for cytotoxic M1 macrophages and *Cd163* (Figure 5d), a marker for M2 macrophages was up-regulated with exercise in the resveratrol treated groups but not the groups on the control diet. Expression of cd86 was however down-regulated with resveratrol treatment in both the sedentary (p<0.001) and exercise (p<0.05) cohorts when compared to the respective control groups.

**Figure 5.**
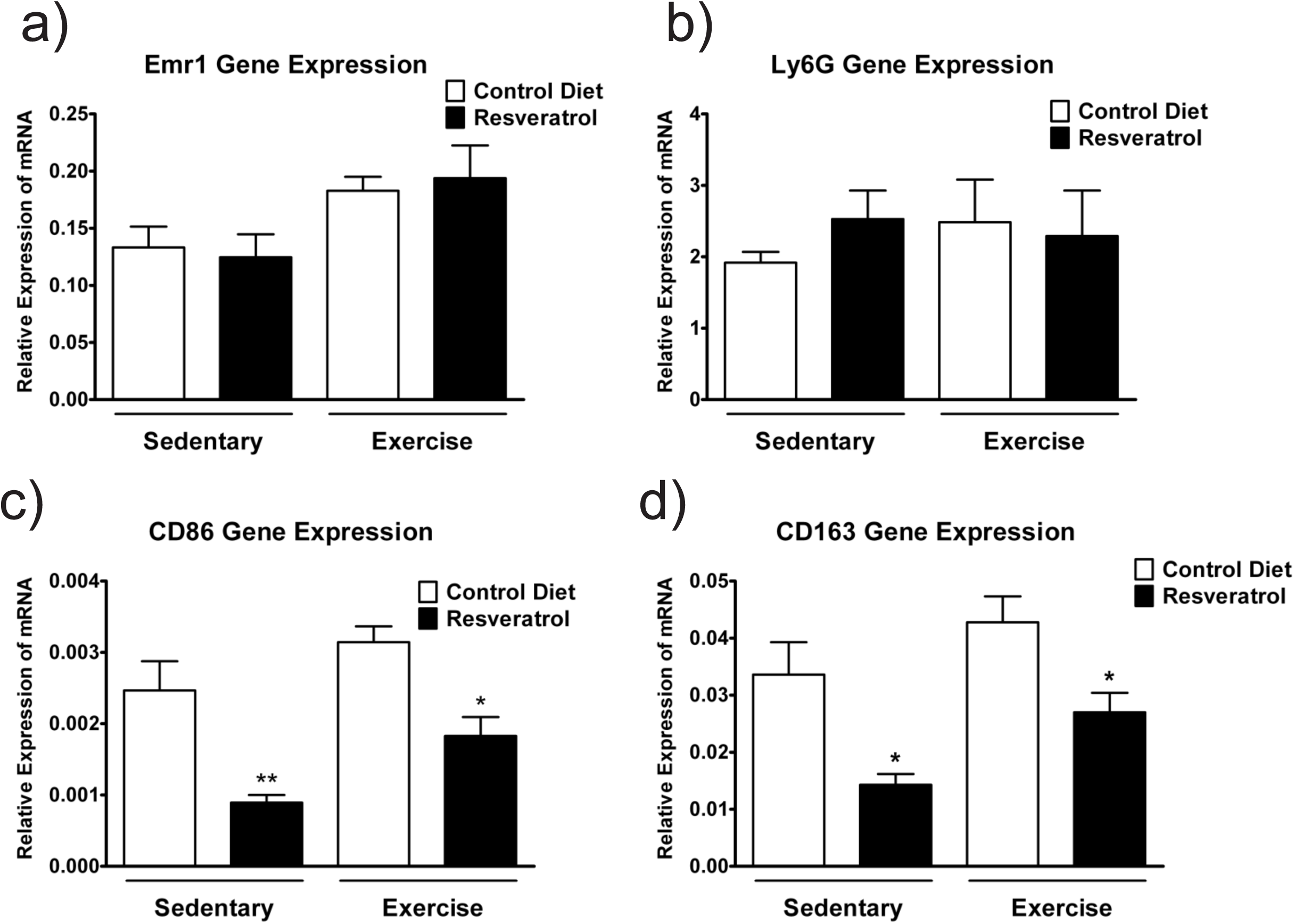
Gene expression of immune cell markers in response to resveratrol administration. a) Expression of Emr1 a marker for (F4/80 positive) macrophages was unchanged with resveratrol treatment in both cohorts. b) Expression of the neutrophil marker Ly6G was unchanged with resveratrol administration in both sedentary and exercise cohorts. c) Expression of CD86, a marker for M1 macrophages, was significantly down-regulated with resveratrol administration in both sedentary and exercise cohorts, when compared to the respective controls. d) Expression of CD163, a marker for M2 macrophages, was significantly down-regulated with resveratrol treatment in sedentary and exercised mice compared to the control cohorts. Graphs show mean ±SEM. qPCR data was analysed via Mann-Whitney U test. Significance is indicated by * p<0.05, ** p<0.001 (n=6).

### Resveratrol treatment does not down-regulate expression of pro-inflammatory cytokines

As gene expression of an M1 macrophage marker was downregulated with resveratrol treatment, we assessed if this was accompanied by a decrease in pro-inflammatory cytokine expression. Interleukin 10 (Il-10) mRNA remained unchanged with resveratrol administration (Figure 6a) in both sedentary and exercise cohorts. Likewise, resveratrol treatment did not change the expression of transforming growth factor beta (TGF-β) in either the sedentary or control groups when compared to the controls (Figure 6d). Interestingly, expression of two major pro-inflammatory cytokines, interleukin 6 (Figure 6b) and tumour necrosis factor (Figure 6c) were significantly increased with resveratrol treatment in the sedentary mice (p<0.05) yet remained unchanged in the exercised mice (Figure 6 c-d).

**Figure 6.**
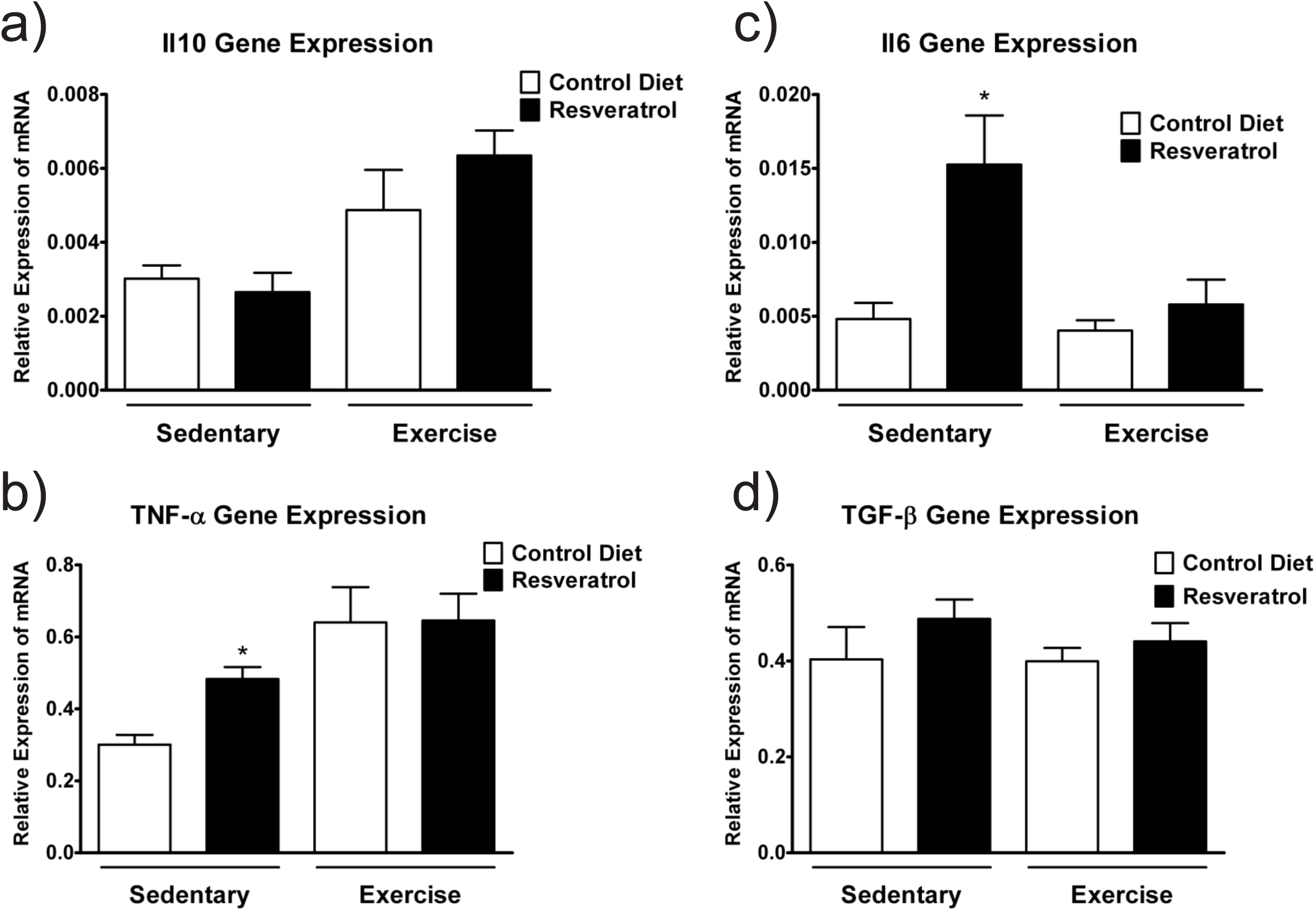
Resveratrol treatment does not down-regulate expression of pro-inflammatory cytokines in *mdx* mice. a) Il-10 gene expression is unchanged with resveratrol treatment in both sedentary and exercise cohorts. b) Il-6 gene expression is significantly up-regulated with resveratrol treatment in the sedentary *mdx* mice but remains unchanged in the exercised group. c) Gene expression of TNF-α is significantly up-regulated with resveratrol treatment in the sedentary *mdx* mice, yet remains unchanged in the exercised group. d) Gene expression of TGF-β is unchanged with resveratrol treatment in both sedentary and exercise groups. Graphs show mean ±SEM. qPCR data was analysed via Mann-Whitney U test. * indicates significance of p<0.05 (n=6).

### Resveratrol treatment does not activate signalling pathways in mdx mice

Previous studies have demonstrated that resveratrol elicits its anti-inflammatory and antioxidant effects through effects on signalling targets such as SIRT1 and NF-κB. SIRT1 belongs to the NAD+-dependent protein deacetylase family or sirtuins which have key roles in many cellular processes including inflammation and oxidative stress. Resveratrol can regulate expression of Sirt1 (Borra et al., 2005; Kaeberlein et al., 2005). The NF-κB pathway is another target of resveratrol and has critical roles in inflammation, immunity, cell proliferation, differentiation and survival (Sadeghi et al., 2017). We assessed if 5mg/kg/day resveratrol was able to alter gene expression of these signalling targets. Sirt1 expression was unchanged with resveratrol administration in both the sedentary and exercise groups when compared to respective controls (Figure 7a). There was no difference in of NF-κB1 (Figure 7b) or NF-κB2 (Figure 7c) expression with the resveratrol treatment in either sedentary or exercised *mdx* mice.

**Figure 7.**
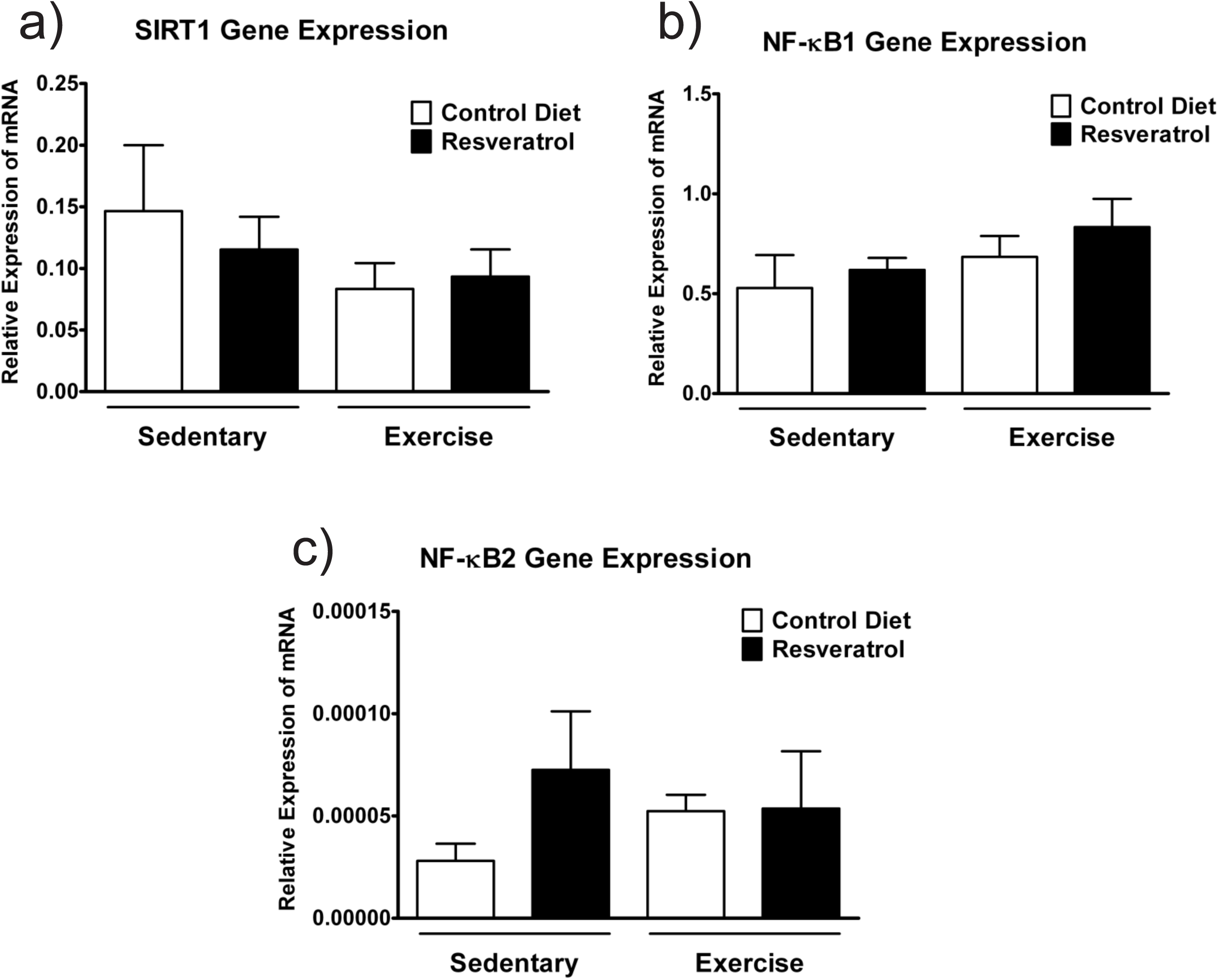
Resveratrol treatment does not alter gene expression of signalling targets. a) Sirt1 gene expression is unchanged with resveratrol treatment in both sedentary and exercise groups when compared to respective controls. b) Gene expression of NF-κB1 is unchanged with resveratrol treatment in both the sedentary and exercise cohorts. c) Likewise, NF-κB2 gene expression is not significantly altered with resveratrol administration in either sedentary or exercised *mdx* mice. Graphs show mean ± SEM. qPCR data was analysed via Mann-Whitney U test (n=6).

## Discussion

### Resveratrol treatment promotes muscle hypertrophy in wildtype mice

To date, most studies that have assessed the effect of resveratrol in myogenic cells have focused on *in vitro* studies in C2C12 cells and primary myoblasts, or have treated *mdx* mice but not analysed the effects in wildtype mice. *In vitro* studies demonstrate that resveratrol administration in C2C12 cells and in primary myoblasts increases myoblast differentiation (Kaminski et al., 2012; Montesano et al., 2013; Saini et al., 2012). Here we show a significant increase in the average myofibre diameter, particularly in fibres which are larger than average, in the *quadriceps* muscle of resveratrol treated wildtype mice. These findings combined suggest that 5mg/kg/day resveratrol treatment promotes muscle hypertrophy *in vivo* and this could have direct applications to the livestock industry or in sports medicine. This effect in wildtype skeletal muscle shows that doses of resveratrol as low as 5mg/kg/day can produce an effect *in vivo.*

### Resveratrol treatment does not improve measures of dystrophic pathology

Treating *mdx* mice with resveratrol did not significantly alter the myofibre diameter. This finding was surprising because hypertrophy was observed in the wildtype skeletal muscle; however, studies in cardiac muscle demonstrate that resveratrol prevents hypertrophy during cardiac stress, such as overload or hypertension (Dolinsky et al., 2013a; Dolinsky et al., 2013b; Thandapilly et al., 2011; Wojciechowski et al., 2010). The dose of resveratrol used in those studies varied but prevention of pressure over-load induced hypertrophy was observed at a dosage of 2.5mg/kg (Wojciechowski et al., 2010). The effect is dependent on the context as in the same study, at the same dosage, volume overload induced hypertrophy was not prevented. As the *mdx* mice are undergoing constant “stress” in the form of inflammation, oxidative stress, muscle degeneration and repair, resveratrol may be acting in a similar way to cardiac muscle and preventing hypertrophy. Alternatively, the 5mg/kg/day dosage used here, while being biologically active, is below the therapeutic threshold in dystrophic muscle.

Resveratrol did not reduce disease parameters in the *mdx* mice. In the sedentary *mdx* mice there was no reduction in the proportion of IgG positive myofibres, skeletal muscle necrosis or serum CK activity with the resveratrol treatment. Again, the lack of efficacy could be attributed to the dosage used here being below the therapeutic threshold. Four published studies have assessed the use of resveratrol in *mdx* mice. In these studies resveratrol administration reduced pathology and improved muscle function (Gordon et al., 2014; Gordon et al., 2013; Selsby et al., 2012) at dosages ranging from 100 to 500 mg/kg/day. Each study was comprehensive in terms of either assessing muscle function (Gordon et al., 2014; Selsby et al., 2012) or downstream signalling effects (Gordon et al., 2013; Hori et al., 2011) however the histological assessment of dystrophic pathology was minimal, with only a single study assessing fibrosis and regenerating myofibres (Hori et al., 2011). As none of these studies assessed skeletal muscle necrosis, damaged myofibres or myofibre diameter, it is difficult to draw parallels between the published studies and the current investigation. Future experiments should consider the effects of the higher concentrations of resveratrol on measures of dystrophic pathology. The resveratrol dose needs to be carefully considered as high mortality rates in mdx mice have been reported at 400mg/kg (Selsby et al., 2012).

While the 5mg/kg bodyweight/day dose of resveratrol did not reduce the percentage of IgG positive fiobres or serm CK levels there was a significant effect of resveratrol on the area of muscle showing signs of necrosis in exercised mdx mice. In mdx mice on a control diet the voluntary exercise intervention used here resulted in a significant increase in damaged tissue in the quadraceps muscle compared to sedentary controls. Of significance In the resveratrol treated mice this increase was not apparent; the area of tissue damage (necrosis) in the quadraceps muscle was similar in sedentary and exercised mice on the resveratrol diet.

### Resveratrol administration reduced gene expression of immune cell markers in dystrophic skeletal muscle but did not reduce expression of inflammatory cytokines

Resveratrol possesses anti-inflammatory properties (Svajger and Jeras, 2012), thus making it an attractive compound for treating the chronic inflammation associated with DMD. Immune cell markers expressed by macrophages and neutrophils were investigated. There are two distinct populations of macrophages which are termed either M1 or M2 macrophages. The M1 “cytotoxic” population release pro-inflammatory cytokines, and can promote cell lysis and phagocytosis of necrotic myofibres, clearing the way for muscle regeneration (Villalta et al., 2009). The M2 anti-inflammatory macrophages release anti-inflammatory cytokines such Il-10 and TGF-β which attenuate the immune response and promote tissue repair (Gordon and Martinez, 2010). Resveratrol treatment significantly down-regulated gene expression of CD86, a marker of cytotoxic M1 macrophages. CD163, a marker of anti-inflammatory M2 macrophages was also significantly downregulated with resveratrol treatment. Expression of the pan-macrophage marker F4/80 (encoded by the Emr1 gene) was unchanged with resveratrol treatment. This was surprising as two of the main macrophage subtypes were significantly down-regulated.

As there was a decrease in the expression of M1 and M2 macrophage markers, the expression of inflammatory cytokines and chemokines was examined. Transcript levels of Il-10 and TGF-β were unchanged with resveratrol treatment in both exercise and sedentary cohorts. Interestingly, levels of IL-6 and TNF-α were increased, rather than decreased, with resveratrol treatment in the sedentary *mdx* mice. These results were not expected, as a decrease in M1 and M2 macrophages would likely be accompanied by decreases in the expression of pro-inflammatory cytokines such as TNF-α. The published higher dosage studies reported similar increases in gene expression with resveratrol treatment in the *mdx* mice, in particular increased expression of Il-6 (Gordon et al., 2013) and TGF-β (Hori et al., 2011). Both studies also showed no significant change in TNF-α gene expression in the resveratrol treated *mdx* mice (Gordon et al., 2013; Hori et al., 2011). These data suggest that resveratrol reduces expression of immune cells markers (Gordon et al., 2013; Hori et al., 2011); however, this effect is not enough to significantly reduce expression of inflammatory cytokines and chemokines such as TNF-α and IL-6.

### Low dose resveratrol does not change gene expression of signalling targets

The anti-inflammatory and anti-apoptotic effects of resveratrol are due to altered signaling pathways, including SIRT1 activation or NF-κB inhibition (Chavez et al., 2008; Howitz et al., 2003; Lagouge et al., 2006). Resveratrol inhibits NF-κB by suppressing the transcriptional activity of P65 (Ren et al., 2013). Due to the role resveratrol has in the activation of SIRT1, Sirt1 gene expression was measured in the quadriceps muscle of the mdx mice. Sirt1 expression was unchanged with the resveratrol treatment in both the sedentary and exercise cohorts. These data are supported by a study which assessed different resveratrol dosages and Sirt1 expression, and concluded that 5mg resveratrol did not significantly up-regulate Sirt1 mRNAand that 100mg was required for Sirt1 up-regulation (Gordon et al., 2013). While mRNA expression is an important measure of SIRT1 some studies show unchanged Sirt1 mRNA with resveratrol treatment but significantly increased activity (Hori et al., 2011). This is important for future studies and shows that Sirt1 gene expression may not be the most reliable measure and should be used in conjunction with measurement of activity. These measurements include commercial ELISA kits which measure NF-κB DNA binding activity (Hori et al., 2011) or through western blotting to measure phosphorylation of the P65 subunit of NF-κB (Ren et al., 2013) in response to resveratrol treatment.

Another signalling target of resveratrol is the NF-κB pathway which has critical roles in inflammation, immunity, cell proliferation, differentiation, and survival. NF-κB and its downstream pro-inflammatory cytokines targets are up-regulated in muscles from DMD patients and in mdx mice (Acharyya et al., 2007; Porreca et al., 1999; Porter et al., 2002). Because NF-κB promotes dystrophic pathology, studies have assessed the impact that reducing NF-κB expression has on dystrophic pathology. A single deletion in the P65 subunit of the NF-κB protein improved muscle morphology (Acharyya et al., 2007; Mourkioti et al., 2006). Resveratrol inhibits NF-κB by suppressing the transcriptional activity of P65 (Ren et al., 2013) and reduces fibrosis in cirrhotic rats by suppressing NF-κB and TGF-β (Chavez et al., 2008). Gene expression of both NF-κB1 and NF-κB2 were not significantly downregulated with the resveratrol treatment in the mdx mice. Like SIRT1, future research would need to assess NF-κB activity in conjunction with gene expression before drawing conclusions about resveratrol’s ability to inhibit NF-κB.

## Conclusions

Overall these experiments produced some interesting findings. Treating healthy wild-type mice with 5mg/kg/day of resveratrol significantly increased myofibre diameter. This finding supports by *in vitro* experiments and could have particular relevance to the livestock or sports medicine fields.

The only improvement observed in the *mdx* mice treated with resveratrol was the prevention of exercise-induced necrosis. This finding is positive as it demonstrates resveratrol can be beneficial in dystrophic muscle and future research can be performed to optimise the concentration at which it is most effective.

Overall, resveratrol has some promising effects in both wildtype and dystrophic skeletal muscle, and future studies to assess multiple concentrations over multiple time-points could determine a resveratrol dosage regime that would provide clinical benefits. Also, the combined administration of resveratrol and corticosteroids could be assessed to determine if there are synergistic therapeutic benefits.

## Acknowledgements

This work was supported by Muscular Dystrophy Australia, Murdoch Children’s Research Institute and the Victorian Government’s Operational Infrastructure Support Program. SRL was supported by a National Health and Medical Research Council of Australia research fellowship [GNT1043837].

## Author Contributions

Keryn Woodman assisted with the design of the study, performed the experiments and wrote the paper. Chantal Coles provided technical advice, helped interpret the data and assisted with writing the paper. Su Toulson maintained the mouse colony. Matthew Knight and Matthew McDonagh did preliminary experiments that led to the study design and provided intellectual assistance. Shireen Lamandé was interpreted the data and revised the paper. Jason White designed the study, interpreted the data, and revised the paper. All authors have read and approved the final manuscript.

